# An economic and robust TMT labeling approach for high throughput proteomic and metaproteomic analysis

**DOI:** 10.1101/2022.07.30.502163

**Authors:** Marybeth Creskey, Leyuan Li, Zhibin Ning, Emily EF Brown, Janice Mayne, Krystal Walker, Anna Ampaw, Robert Ben, Xu Zhang, Daniel Figeys

**Affiliations:** Regulatory Research Division, Centre for Oncology, Radiopharmaceuticals and Research, Biologic and Radiopharmaceutical Drugs Directorate, Health Products and Food Branch, Health Canada, Ottawa, Canada; School of Pharmaceutical Sciences, Ottawa Institute of Systems Biology and Department of Biochemistry, Microbiology and Immunology, Faculty of Medicine, University of Ottawa, Ottawa, Canada; Department of Chemistry, Faculty of Science, University of Ottawa, Ottawa, Canada

**Keywords:** high throughput, metaproteomics, microbiome, robust, TMT workflow, proteomics

## Abstract

Multiplexed quantitative proteomics using tandem mass tag (TMT) is increasingly used in –omic study of complex samples. While TMT-based proteomics has the advantages of the higher quantitative accuracy, fewer missing values, and reduced instrument analysis time, it is limited by the increased cost due to the use of labeling reagents. In addition, current TMT labeling workflows involve repeated small volume pipetting of reagents in volatile organic solvents, which may increase the sample-to-sample variations and is not readily suitable for high throughput applications. In this study, we demonstrated that the TMT labeling procedures could be streamlined by using pre-aliquoted dry TMT reagents in a 96 well plate or 12-tube strip. As little as 50 μg dry TMT per channel effectively labels 6-12 μg peptides, yielding efficient TMT labeling efficiency (∼99%) in both microbiome and mammalian cell line samples. This streamlined workflow decreases reagent loss and reduces inter-sample variations. We applied this workflow to analyze 97 samples in a study to evaluate whether ice recrystallization inhibitors improve the cultivability and activity of frozen microbiota. The results demonstrated tight sample clustering corresponding to groups and consistent microbiome responses to prebiotic treatments. This study supports the use of TMT reagents that are pre-aliquoted, dried, and stored for streamlined and robust quantitative proteomics and metaproteomics in high throughput applications.

## Introduction

High-resolution mass spectrometry (MS)-based proteomic and metaproteomic technologies allow for deep insights into protein- and proteoform-level information of many types of samples, including cells, tissues and microbiomes. Along with other –omic approaches, proteomics has been widely applied for large-scale clinical sample analysis [1-3], enabling extensive molecular profiling and biomarker discovery for human health management. Proteomics and metaproteomics have also been applied in *in vitro* or *ex vivo* assays [4], allowing for proteomics-based high content, high throughput drug screenings. For the study of microbiome, metaproteomics not only provides quantitative information of expressed microbial proteins that estimate the functional activities, but also can be used to assess the microbiome biomass and analyze community structures [4, 5]. These advantages make proteomics and metaproteomics suitable for screening and evaluation of drugs that target host-microbiome interactions. The latter are increasingly implicated in various diseases ranging from gastrointestinal, metabolic diseases to neurological disorders and cancers [6-12].

The development of multiplexing approaches, such as isobaric tandem mass tags (TMT), dramatically reduces the MS time required per sample, which vastly improves the throughput of proteomic analyses. Currently, by using the TMT approach, up to 18 samples can be multiplexed [13]. However, the application of TMT-based proteomics and metaproteomics approaches in high throughput sample analysis increases the reagent cost and introduces additional steps during sample preparation. In a previous version of the manufacturer’s instruction, 800 μg TMT10 reagent per sample (corresponding to ∼120 US dollars) in 141 μl reaction volume was recommended to label 25 -100 μg peptides. More recently, new versions of TMT10 reagents (catalog 90308 and 90309) are packed as 200 μg aliquots that are recommended to label 10 - 25 μg protein digests in 45 μl reaction volume, which reduces the cost to ∼30 US dollars per sample. Zecha *et al*. reported that the TMT-to-peptide ratio could be markedly reduced from 8: 1 to 1: 1 while maintaining sufficient labeling efficiency [14]. Additionally, they demonstrated that the TMT labeling reaction volume can be decreased to 25 μl with 100 μg TMT reagent, which further reduced the cost of TMT reagents per sample (to ∼15 US dollars). While the reaction volume was decreased, the TMT labeling workflow remained the same, which involved pipetting of TMT reagents in small volumes of volatile organic solvents (i.e., acetonitrile; 5 - 20 μl). Transferring small volumes of acetonitrile (ACN) in a high throughput manner is challenging, which makes the conventional TMT labeling workflow difficult for high throughput applications.

In this study, we established a streamlined robust TMT labeling procedure using 96-well plates or 12-strip tubes pre-loaded with dry TMT reagents. In this streamlined TMT labeling workflow, the protein digest was re-suspended in sample buffer containing 20 - 30% (v/v) ACN and directly added to the dry TMT plate or tube for labeling. We demonstrated that as little as 50 μg TMT reagent in 15 μl reaction volume was sufficient to efficiently label 6 - 12 μg peptides and the workflow was readily compatible with high throughput sample processing. With reduced inter-sample variations and increased reagent utilization efficiency (due to the large-batch preparation of TMT plates and tube-strips), the established workflow provides an economic (∼ 7.5 US dollars per sample), robust, and easy-to-implement workflow for multiplexed quantitative proteomics and metaproteomics, enabling applications in various fields, such as high throughput drug screening and large-scale clinical studies.

## Experimental Section

### Sample collection and processing for Caco-2 cells and human microbiota

Human intestinal Caco-2 cells were purchased from the American Type Culture Collection (ATCC) and grown at in Dulbecco's Modified Eagle Medium (DMEM) supplemented with 10% fetal bovine serum at 37°C with 5% CO_2_. Human stool samples were collected from healthy volunteers with a protocol approved by the Ottawa Health Science Network Research Ethics Board at the Ottawa Hospital (Ottawa, Canada). Protocols for stool collection, pre-processing, live microbiota bio-banking, and *ex vivo* microbiome culturing were described previously [4, 15].

Protein extractions and trypsin digestion from microbiota and Caco-2 cells were performed using different protocols to demonstrate the wide applicability of the established dry TMT workflow. Briefly, microbiota cells were lysed in 4% SDS (w/v), 8 M urea, 50 mM Tris-HCl (pH 8.0) buffer. SDS was then removed with precipitation using 5-fold volumes of ice-cold acetone/ethanol/acetic acid (50/50/0.1 (v/v/v)) buffer overnight at −20°C. Proteins were pelleted, washed with acetone, and re-suspended in 6 M urea, 50 mM ammonium bicarbonate for reduction with dithiothreitol (DTT), alkylation with iodoacetamide (IAA), in-solution trypsin digestion, and desalting. The resulting peptides were dried and re-suspended in 100 mM triethylamonium bicarbonat (TEAB) for TMT labeling. Caco-2 cells were lysed in 1% SDS, 100 mM TEAB buffer. Proteins were reduced with tris(2-carboxyethyl)phosphine (TCEP) then alkylated with IAA. Protein was then precipitated with the addition of 6 volumes of ice-cold acetone. The protein was pelleted, air-dried briefly, dissolved to 2 μg/μl in 100 mM TEAB followed by trypsin digestion. After digestion, the peptide solution was diluted in 100mM TEAB and used directly for TMT labeling. More details on microbiota and Caco-2 cell culturing and protein/peptide sample preparation are shown in Supporting Information.

### TMT reagents preparation and labeling

For the evaluation of TMT labeling efficiency, equal amount of the 11 channels of TMT11plex label reagents (catalog number A34808, ThermoFisher Scientific Inc.) were pooled to generate a TMT mix. A portion of the TMT mix was stored in ACN solution at −80°C until use, and the remaining was aliquoted into low-binding tubes with 50 μg or 100 μg per aliquot, completely dried using a vacuum concentrator. Conventional TMT labeling workflow was performed by mixing wet TMT reagent (in ACN) with peptide solution dissolved in 100 mM TEAB. For dry TMT labeling, peptides were dissolved or diluted in 100 mM TEAB (with 20-30% ACN) and added directly into the tubes containing dry TMT reagents. All reactions were carried out at 25°C for 2 hours, and quenched using a final concentration of 0.4% hydroxylamine for 15 min.

To prepare dry TMT plates or tube strips, 5 mg TMT11plex label reagents were re-suspended in pure ACN to make 1 mg/ml stock solution, and aliquoted into plates or strip tubes with 50 μg reagent in each well/tube. The reagent plates/tube strips were then completely dried using a vacuum concentrator and stored at −80°C prior to use. For convenience, a 96-well plate and a 12-tube strip were used with each row or strip being a full labeling reagent set as indicated in Figure 1 (left to right: 126, 127N, 127C, 128N, 128C, 129N, 129C, 130N, 130C, 131N, 131C, empty). For peptide labeling, 15 μl of peptide solution was added directly to each tube or well, incubate at 25°C for 2 hours, and quenched with hydroxylamine. The individual labels were then combined to generate a mixture for either desalting and LC-MSMS analysis or fractionation using high pH reverse-phase chromatography prior to LC-MSMS analysis. More details on TMT labeling procedures are shown in Supporting Information.

**Figure 1.**
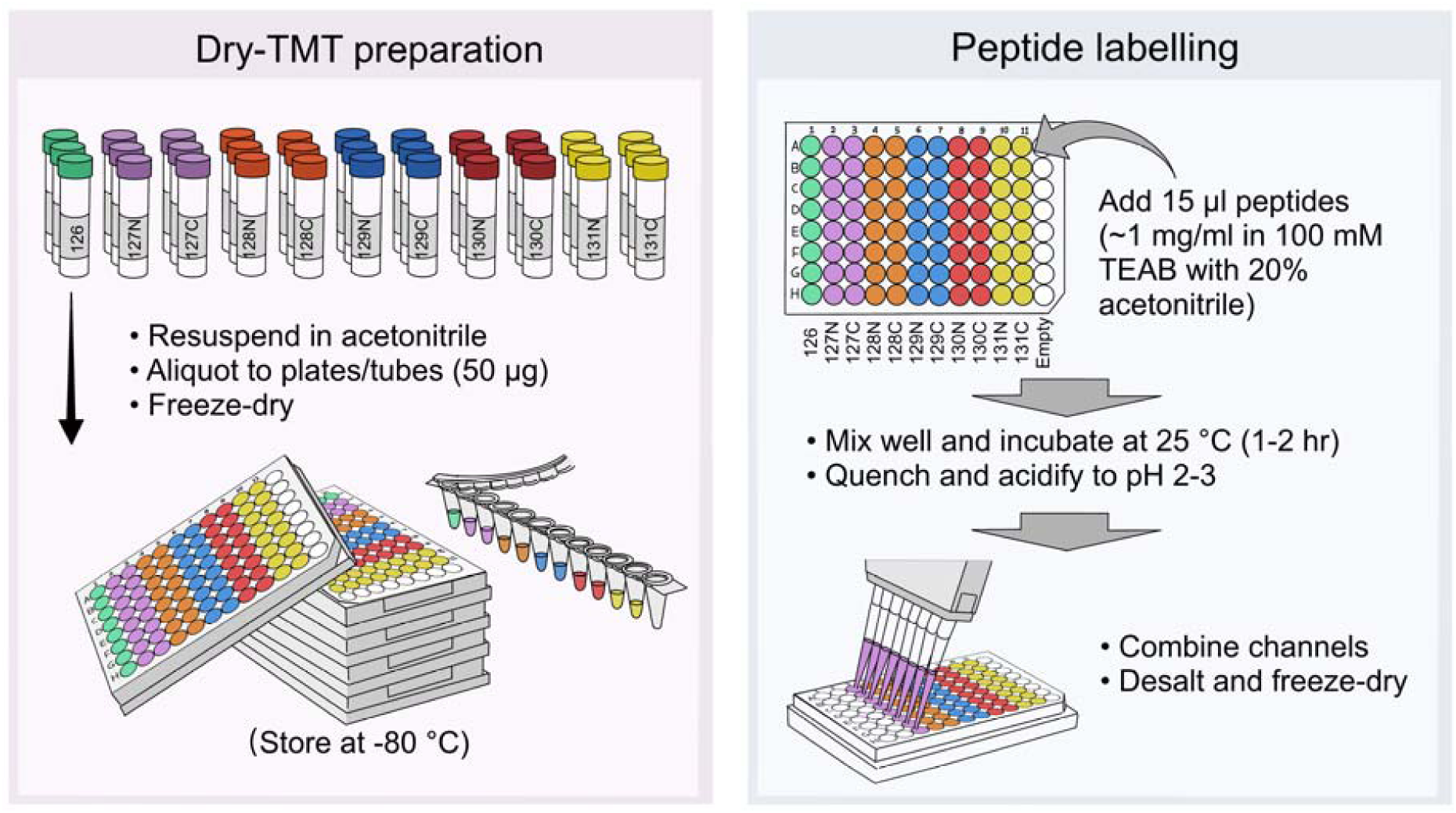
Workflow of dry TMT plate- or tube strip- based TMT labeling for high throughput quantitative proteomic and metaproteomic sample analyses.

### *LC-MSMS analysis* and bioinformatics

Tryptic peptides derived from microbiome and Caco-2 cells were analyzed using Thermo Scientific(tm) Orbitrap Exploris 480 and Fusion Lumos Tribrid mass spectrometers, respectively. More details on instrument setup and parameters are shown in Supporting Information. All data were recorded by Xcalibur and exported into RAW format for bioinformatics processing.

The database searches for the gut microbiome data set were performed using MetaLab (version 2.1) with an integrated human gut microbial gene catalog (IGC) database (containing 9.9 million microbial protein sequences) and MetaPro-IQ workflow [16-18]. The database searches of Caco-2 data set were performed using both MaxQuant and Proteome Discoverer 2.4 with a human SwissProt database (downloaded March 9, 2022, 20307 entries). For TMT labeling efficiency evaluation, TMT (+ 229.163 Da) was specified as variable modifications on lysine and peptide N termini; and for over-labeling evaluation, database search was performed with TMT mode and, additionally, TMT on histidine (H) or serine (S), threonine (T), and tyrosine (Y) were set as variable modifications. For quantitation, the database search was performed with a standard TMT mode. For all searches, Carbamidomethyl (C) was set as a fixed modification, and Oxidation (M) and Acetyl (Protein N-term) as variable modifications.

All identified PSMs were used for the calculation of TMT labeling efficiency. Only PSMs with TMT modification on all lysine resides and free peptide N termini were considered as “fully labeled”. PSMs that did not have any TMT modification and has at least one accessible amine were considered as “not labeled”. PSMs that contained at least one TMT modification and were not fully labeled were considered as “partially labeled”. Over-labeling rate was calculated by dividing the number of identified PSMs with TMT modification on either H, S, T or Y by the total number of identified PSMs. More details on the bioinformatics procedures, labeling efficiency calculation and plotting are shown in Supporting Information.

### Data availability

All MS proteomics data along with the database search results were deposited to the ProteomeXchange Consortium (http://www.proteomexchange.org) via the PRIDE partner repository.

## Results and Discussion

In conventional TMT labeling workflow, TMT reagents are equilibrated to room temperature, re-suspended in ACN and used immediately for peptide labeling (leftover reagent suspension can be stored at −20 °C or −80 °C for around 3 months prior to usage). This is challenging for high throughput applications where small reaction volumes are usually used to reduce the reagent cost. Currently, as little as 25 μl reaction volume has been applied for TMT labeling with 5 μl (100 μg) of TMT reagents in ACN mixed with 20 μl peptide samples [14]. Extra caution is needed for accurate transfer of 5 μl ACN solution given its nature of high volatility, which otherwise may result in poor inter-sample consistency and impact the robustness of the quantitation. To streamline the TMT labeling workflow in a manner that is suitable for high throughput applications, in this study TMT reagents were aliquoted into low-binding 96-well plates or 12-strip tubes with as little as 50 μg reagents (in 50 μl ACN) in each well or tube (Figure 1). While even smaller amount of TMT per channel is possible, to be compatible with high throughput applications using automatic liquid handing system, this study only tested 50 μg and 100 μg TMT per channel that require 15 μl and 25 μl of reaction volume, respectively. The TMT plates or strip tubes were then dried and stored at −80 °C prior to peptide labeling. In this study, we selected TMT11plex to fit into a row (12 wells) of a 96-well plate or a 12-tube strip with the first well/tube being loaded with TMT channel 1 (TMT11-126) and sequentially loaded until the 11^th^ well/tube with channel 11 (TMT11-131C) of TMT reagents (Figure 1). For TMT labeling, ∼1 mg/ml peptide solutions in 100 mM TEAB containing 20 - 30% (v/v) ACN were prepared and 15 μl of them were added directly to the dried TMT plate or tube for incubation at room temperature for 1 - 2 hours (Figure 1). The individually labeled samples were then quenched and pooled row-wise followed by high pH reverse-phase fractionation or direct desalting and MS analysis.

In addition to the streamlined application for high throughput sample processing, the large-batch preparation of dry TMT reagent plates or tube strips holds the promise to greatly reduce reagent loss and enable better sample-to-sample consistency thanks to the relatively larger volumes of pipetting (TMT reagents can be diluted to ≤ 1 μg/μl). To evaluate whether the use of dry TMT reagents achieves sufficient TMT labeling efficiency, we compared the dry TMT workflow with the conventional labeling workflow (Wet TMT) in two different sample types (microbiome and intestinal Caco-2 cells) with two different instrument platforms (Orbitrap Exploris 480 and Orbitrap Fusion Lumos). Briefly, we equally mixed 11 channels of TMT11plex to prepare aliquots of 100 μg wet TMT mix in 5 μl ACN (group W100), 100 μg dry TMT mix (group D100), or 50 μg dry TMT mix (group D50). For microbiome samples, two different amounts of peptides were used for labeling for each group with 25 μl reaction volume for 100 μg TMT groups and 15 μl reaction volume for 50 μg TMT group (Figure 2A). Metaproteomic database searches were performed using MetaLab with IGC as a database and TMT11 (lysine and peptide N termini) as variable modifications. An average of 7,307 ± 260 peptide-spectrum matches (PSMs) per sample, corresponding to 7,206 ± 235 peptides, were identified and there was no obvious difference of identification between groups (Figure 2A). TMT full labeling efficiency of ∼ 99% was obtained for all samples (98.7% - 99.6%) and we did not observe obvious difference between Wet TMT and Dry TMT groups with both high and low peptide inputs (Figure 2B). A TMT labeling efficiency of >95% is considered acceptable for accurate quantifications [19, 20], and therefore the established dry TMT workflow maintained efficient TMT labeling efficiency. In this study, we further increased the depth of measurement with 120 min MS run times, which did not change the findings (Figure 2D-E). In the evaluations using Caco-2 cell lysate samples, consistent and similar TMT labeling efficiencies for both 50 μg and 100 μg TMT groups were obtained (>99% for all samples; Figure 2G-H). Similar observations were obtained when the Caco-2 dataset was searched using the Proteome Discoverer (Figure S1). Evaluation of the over labeling rate on non-lysine amino acid residues (i.e., H, S, T, Y) showed no obvious difference between groups in both microbiome and Caco-2 cell lysate samples with < 5% over labeling rate for all tested samples (Figure 2C, F and I). This was at a similar level with the findings in a previous study [14].

**Figure 2.**
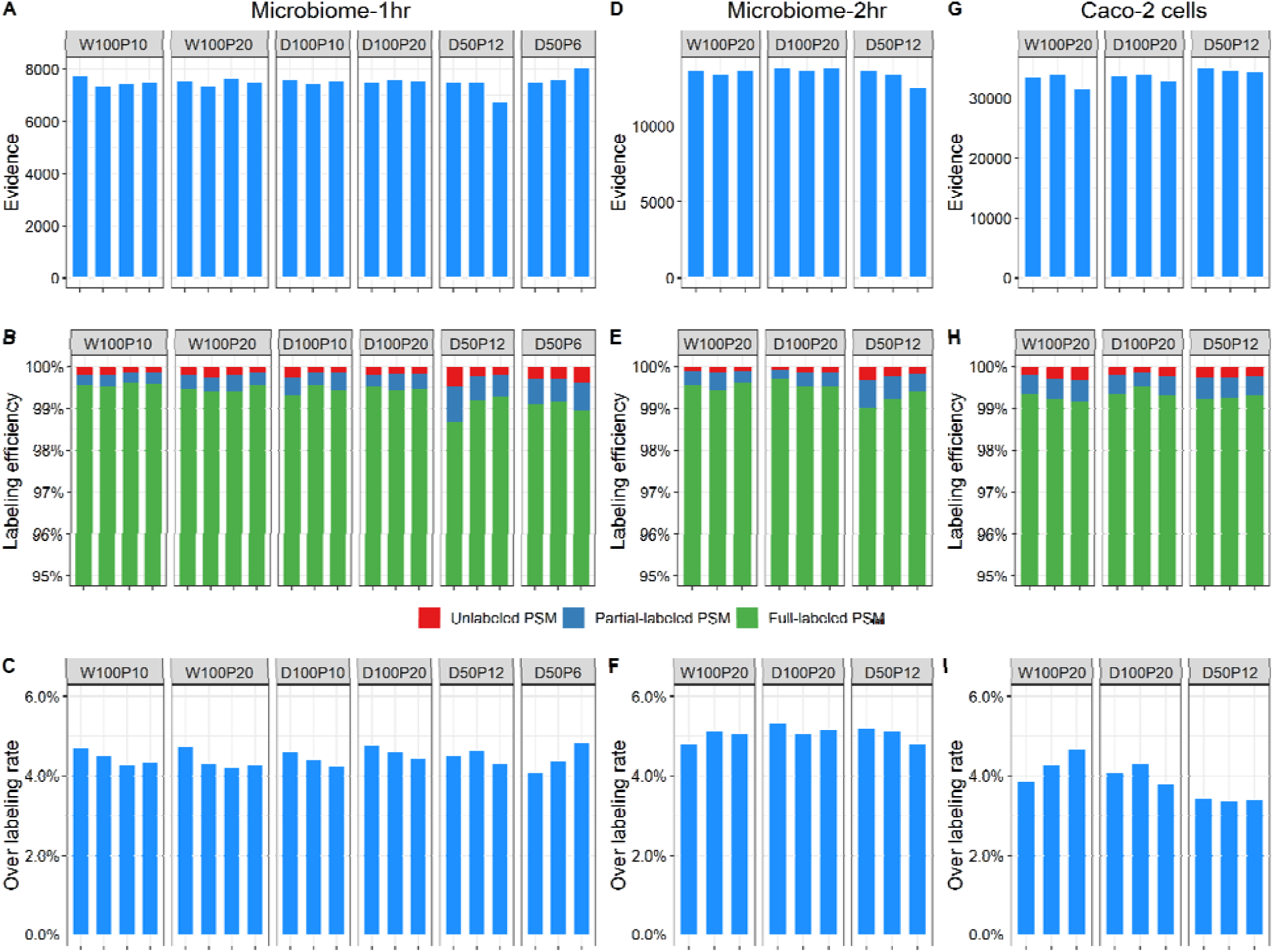
Comparison of labeling efficiency between convenient dry TMT labeling workflow with standard wet TMT workflow. (A-C) Identified PSM evidence (upper), TMT labeling efficiency (middle), and over labeling rate (lower) in microbiome samples with 1 hr MS run; (D-F) Identified PSM evidence (upper), TMT labeling efficiency (middle), and over labeling rate (lower) in microbiome samples with 2 hr MS run; (G-I) Identified PSM evidence (upper), TMT labeling efficiency (middle), and over labeling rate (lower) in Caco-2 cell samples. Each bar in the plots represents one replicate in the group.

We then evaluated the quantitative accuracy of the established dry TMT labeling workflow with the analysis of 22 same aliquots of Caco-2 cell lysate samples. Briefly, twenty-two 12 μg peptide aliquots derived from Caco-2 cells were labeled using two strips of dry TMT reagents (50 μg TMT per channel), generating two mixtures (Strip 1 and Strip 2). Both single-shot (90 min) and fractionated (90 min of 8 fractions) MS analyses were performed to assess whether similar observations can be obtained using deep measurements. In agreement with the above evaluations using pooled TMT reagents, >99% labeling efficiency was achieved for unfractionated single-shot sample analysis and, as expected, slightly lower TMT labeling rate was observed for fractionated samples (98.4% and 98.7% for Strip 1 and Strip 2, respectively) (Figure S2). Nevertheless, these TMT labeling efficiencies in fractionated samples were still higher than 95% that is considered acceptable for accurate quantifications [19, 20]. In this experiment, MaxQuant search quantified 2093 and 4690 protein groups, corresponding to 10,101 and 37,747 peptide sequences, for single-shot and fractionated samples, respectively. Abundance normalization was then performed using MSstatsTMT workflow [21], and the normalized intensities of protein groups were used to calculate the log2-transformed ratios to reference channel (channel 1) for each protein group in each sample. As shown in Figure 3, the calculated abundance ratios were consistently distributed and close to one, the theoretical value (i.e., log2-transformed ratio of 0 in the plot), for both fractionated and single-shot samples. These observations indicate high quantitative accuracy using the dry TMT labeling workflow.

**Figure 3.**
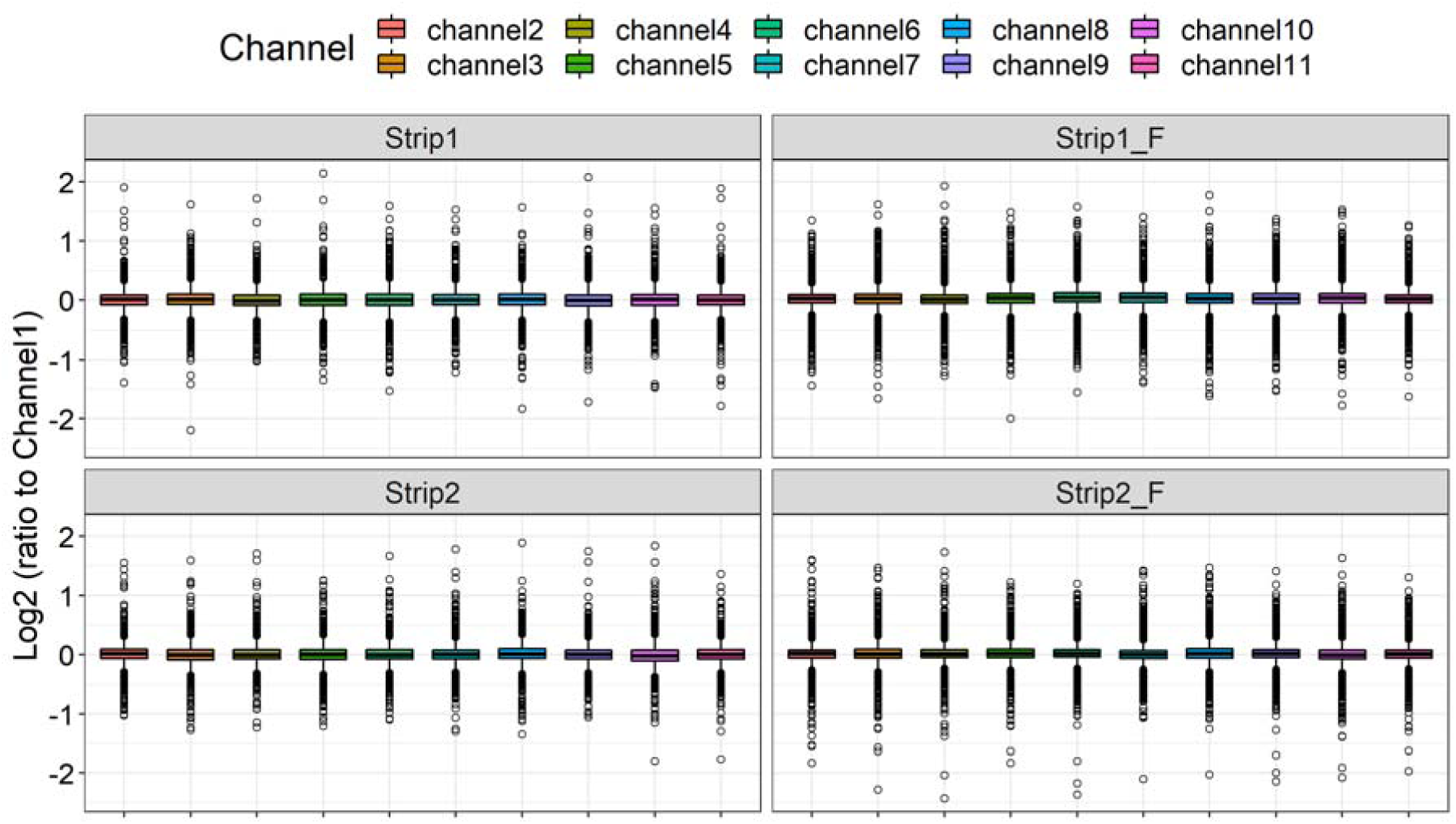
Evaluating the quantitative accuracy for Caco-2 cell lysate samples. The same samples were labeled using strip tubes with 50 μg dry TMT reagent each tube. Mixtures were either fractionated (right panel) or not (left panel) prior to MS analysis. Ratios to reference channel (channel 1) of all quantified protein groups were calculated, log2-transformed, and used for plotting in R using *ggboxplot* function.

To further evaluate the quantitation of the dry TMT workflow, we applied this approach in a study to evaluate whether different ice recrystallization inhibitors (IRIs) improve the cultivability and treatment-responses of frozen microbiota in a high throughput *ex vivo* microbiome assay. Briefly, fresh stool collected from a healthy adult volunteer was processed and stored at −80 °C for two weeks with or without two different IRIs, i.e., β-4-bromophenyl-D-glucose (IRIa) and *N*-2-fluorophenyl-D-gluconamide (IRIb) [22, 23], at five different concentrations of each. Both fresh stools and the frozen microbiotas were cultured *in vitro* with and without 5 mg/ml kestose, a prebiotic with known effects on modulating the microbiota, for 24 hours using a previously established RapidAIM assay [4]. In total, 97 samples were generated and processed for shotgun metaproteomic analysis using dry TMT labeling workflow. A small aliquot of each sample was combined to generate a pooled sample, which was used as reference channel for each TMT experiment. We randomized the 97 samples into two dry TMT 96-well plates (Plate1 row A-H and Plate2 row A-B, labeled as P1A-H and P2A-B, respectively) with the channel 1 of each row as reference (pooled sample) (the remaining 3 wells were added with pooled samples as well; Figure S3). Three additional rows (i.e., 33 wells; Plate2 row C-E, labeled as P2C, P2D and P2E, respectively) were used to label pooled sample for all channels to evaluate the inter-channel variations. Metaproteomic analysis of the resulting 13 mixtures using 2-hour single-shot MS run identified 28,605 microbial peptides and 10,656 protein groups. In agreement with the findings using Caco-2, the calculated abundance ratios were consistently distributed and close to one for the pooled sample experiments (P2C, P2D and P2E) (Figure 4). On the contrary, the ratio distributions of the IRI study samples (P1A to P1H and P2A to P2B) showed marked higher variations than pooled sample experiments (Figure 4), suggesting marked changes of protein expressions in response to either treatments or storage conditions of the frozen microbiotas.

**Figure 4.**
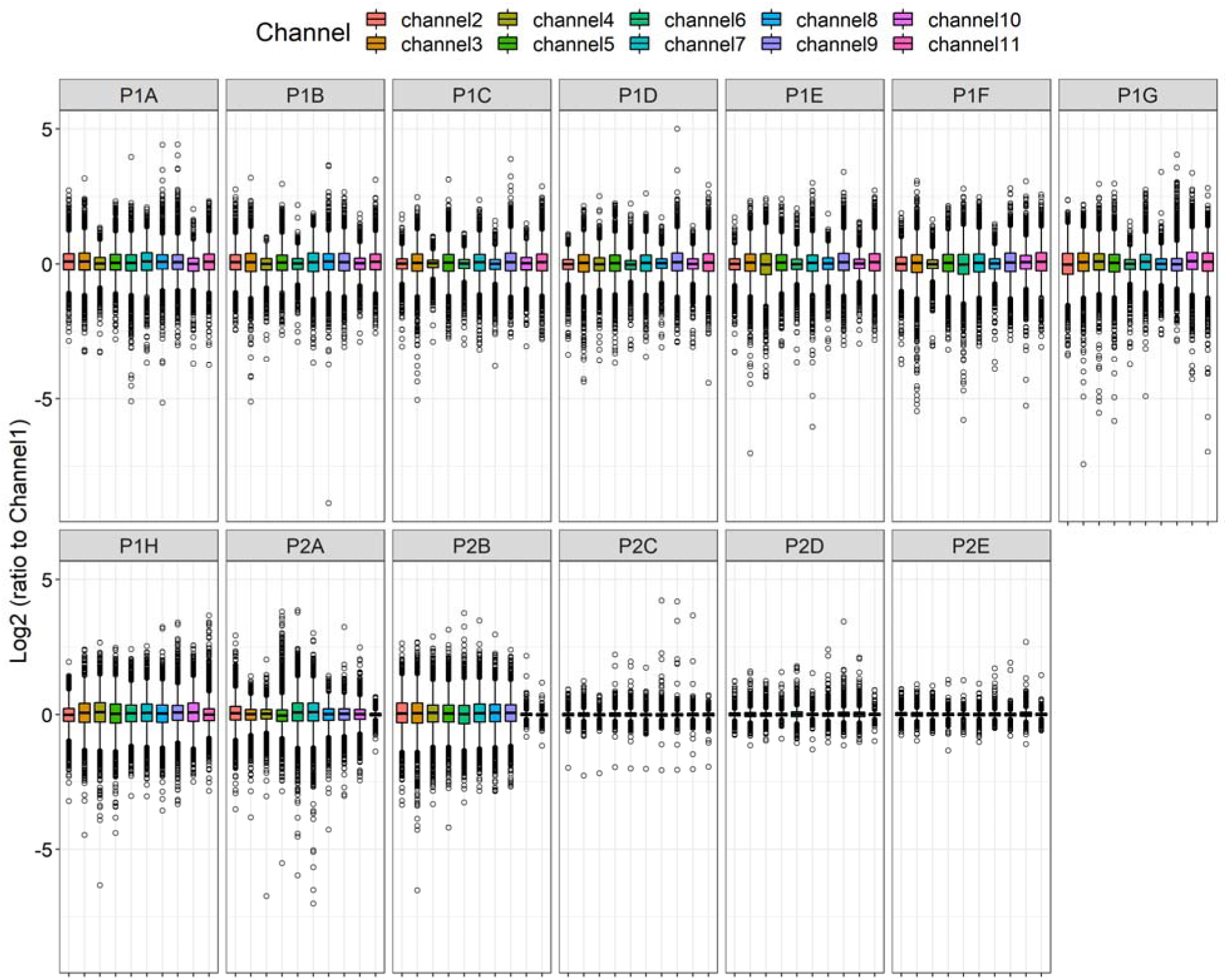
Distribution of the quantified ratios of all quantified protein groups in the IRI microbiome study. Ratios of protein intensity to the corresponding proteins in reference channel (channel 1) in each mixture were calculated and log2-transformed for the plotting in R using *ggboxplot* function.

To evaluate the influences of kestose treatment and IRI supplement on microbiome, unsupervised principal component analysis (PCA) was performed using the abundance ratios of quantified gut microbial protein groups. As shown in Figure 5, clear sample clustering according to either the treatment or the different inoculating microbiotas was observed. Baseline (BL) group samples (microbiotas prior to *ex vivo* culturing; plotted as squares) clustered together at the top along the second principal component (vertical axis) and accounted for 18% of the variations. Marked differences between the metaproteomes of cultured microbiomes with and without the treatment of kestose were observed along the first principal component (horizontal axis), which explains 49% of the variations displayed in PCA score plot. These results are in agreement with previous studies showing the modulating effects of kestose on *in vitro* cultured microbiota [24, 25]. As expected, microbiome culturing using fresh stools clustered closer to the baseline samples compared to those using frozen microbiotas. No obvious difference was observed with the addition of IRIb for freezing microbiota, while IRIa influenced the metaproteomic profiles of the cultured microbiome in a dose-dependent manner in the kestose groups (Figure 5). These observations may indicate that micro-organisms within the microbiome are capable of metabolizing IRIa (a glucose derivate), which needs further investigation to better demonstrate its suitability for use in live microbiota bio-banking.

**Figure 5.**
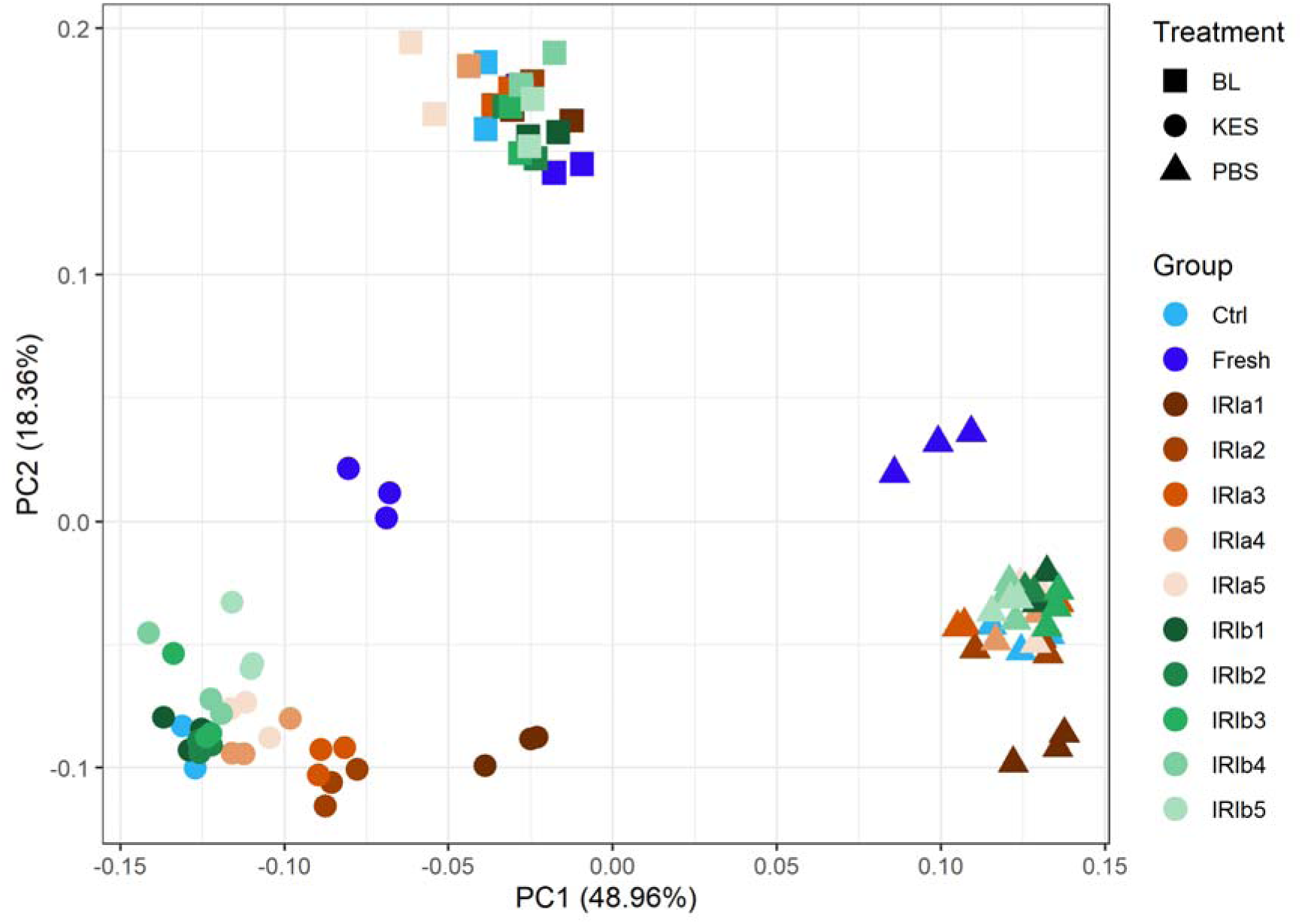
PCA score plot of *ex vivo* cultured metaproteomes with or without kestose and IRIs. The first two principal components (PC1 and PC2) were displayed. BL, baseline; KES, kestose treatment at 5 mg/ml; PBS, phosphate buffered saline as vehicle control; Ctrl, control microbiome frozen without IRI; Fresh, fresh microbiome for culturing; IRIa 1 - 5 and IRIb1 - 5, different concentrations of IRIa or IRIb with 1 as the highest and 5 as the lowest concentrations.

## Conclusions

In this study, we established a TMT labeling workflow that uses pre-aliquoted dry TMT reagents in 96 well plates or 12-tube strips, which streamlines the TMT labeling procedure, reduces the cost of experiments, and improves the inter-sample or inter-batch consistency for proteomic quantitation. Evaluations of the labeling efficiency using human gut microbiome and Caco-2 cell samples both demonstrated efficient TMT labeling rate (∼99%) and no increased deleterious off-target labeling. This study was performed using TMT11plex reagent, however, the workflow and implementation is readily applicable to other TMT formats, including TMTpro (up to 18plex). We showed that as little as 50 μg TMT reagent was able to sufficiently label up to 12 μg peptides in a 15 μl reaction volume. It is possible that the amount of reagents being used, TMT-to-peptide ratio, and/or reaction volume can be further reduced [14], however careful attention needs to be taken to sufficiently mix the TMT reagents with peptides to obtain optimal labeling efficiency. An outstanding advantage of this workflow is that there is no need to pipette TMT reagents in organic solvents (i.e., acetonitrile) during sample labeling, which makes the established TMT labeling workflow a perfect fit for high throughput applications, such as drug screening and large-scale clinical sample analysis, with automated liquid handling system.

## Supporting Information

The Supporting Information is available online at https://pubs.acs.org/.

Detailed descriptions on experimental procedures (DOC); TMT labeling efficiency in Caco-2 cell samples using Proteome Discoverer database search (Figure S1); TMT labeling efficiency of fractionated and unfractionated Caco-2 samples (Figure S2); Layout of IRI study samples in 96-well TMT plates (Figure S3).

## Acknowledgement

This work was supported by the Government of Canada through Health Canada, Genome Canada and the Ontario Genomics Institute [OGI-156 and OGI-149], the Natural Sciences and Engineering Research Council of Canada [NSERC, grant no. 210034], and the Ontario Ministry of Economic Development and Innovation [ORF-DIG-14405 and project 13440]. We also gratefully acknowledge Dr. Simon Sauvé, Dr. Daryl Smith and Dr. Roger Tam from Health Canada for commenting and editing on the manuscript.

## References

1. Tebani, A., et al., Integration of molecular profiles in a longitudinal wellness profiling cohort. Nat Commun, 2020. 11(1): p. 4487.

2. Chen, R., et al., Personal omics profiling reveals dynamic molecular and medical phenotypes. Cell, 2012. 148(6): p. 1293–307.

3. Messner, C.B., et al., Ultra-High-Throughput Clinical Proteomics Reveals Classifiers of COVID-19 Infection. Cell Syst, 2020. 11(1): p. 11–24 e4.

4. Li, L., et al., RapidAIM: a culture- and metaproteomics-based Rapid Assay of Individual Microbiome responses to drugs. Microbiome, 2020. 8(1): p. 33.

5. Kleiner, M., et al., Assessing species biomass contributions in microbial communities via metaproteomics. Nat Commun, 2017. 8(1): p. 1558.

6. Cho, I. and M.J. Blaser, The human microbiome: at the interface of health and disease. Nat Rev Genet, 2012. 13(4): p. 260–70.

7. Gilbert, J.A., et al., Current understanding of the human microbiome. Nat Med, 2018. 24(4): p. 392–400.

8. Baruch, E.N., et al., Fecal microbiota transplant promotes response in immunotherapy-refractory melanoma patients. Science, 2021. 371(6529): p. 602–609.

9. Davar, D., et al., Fecal microbiota transplant overcomes resistance to anti-PD-1 therapy in melanoma patients. Science, 2021. 371(6529): p. 595–602.

10. Gopalakrishnan, V., et al., Gut microbiome modulates response to anti-PD-1 immunotherapy in melanoma patients. Science, 2018. 359(6371): p. 97–103.

11. Matson, V., et al., The commensal microbiome is associated with anti-PD-1 efficacy in metastatic melanoma patients. Science, 2018. 359(6371): p. 104–108.

12. Routy, B., et al., Gut microbiome influences efficacy of PD-1-based immunotherapy against epithelial tumors. Science, 2018. 359(6371): p. 91–97.

13. Li, J., et al., TMTpro-18plex: The Expanded and Complete Set of TMTpro Reagents for Sample Multiplexing. J Proteome Res, 2021. 20(5): p. 2964–2972.

14. Zecha, J., et al., TMT Labeling for the Masses: A Robust and Cost-efficient, In-solution Labeling Approach. Mol Cell Proteomics, 2019. 18(7): p. 1468–1478.

15. Zhang, X., et al., Evaluating live microbiota biobanking using an ex vivo microbiome assay and metaproteomics. Gut Microbes, 2022. 14(1): p. 2035658.

16. Zhang, X., et al., MetaPro-IQ: a universal metaproteomic approach to studying human and mouse gut microbiota. Microbiome, 2016. 4(1): p. 31.

17. Cheng, K., et al., MetaLab: an automated pipeline for metaproteomic data analysis. Microbiome, 2017. 5(1): p. 157.

18. Li, L., et al., iMetaLab Suite: A one-stop toolset for metaproteomics. iMeta, 2022: p. e25.

19. Erdjument-Bromage, H., F.K. Huang, and T.A. Neubert, Sample Preparation for Relative Quantitation of Proteins Using Tandem Mass Tags (TMT) and Mass Spectrometry (MS). Methods Mol Biol, 2018. 1741: p. 135–149.

20. Hogrebe, A., et al., Benchmarking common quantification strategies for large-scale phosphoproteomics. Nat Commun, 2018. 9(1): p. 1045.

21. Huang, T., et al., MSstatsTMT: Statistical Detection of Differentially Abundant Proteins in Experiments with Isobaric Labeling and Multiple Mixtures. Mol Cell Proteomics, 2020. 19(10): p. 1706–1723.

22. Jahan, S., et al., Inhibition of ice recrystallization during cryopreservation of cord blood grafts improves platelet engraftment. Transfusion, 2020. 60(4): p. 769–778.

23. Poisson, J.S., et al., Modulating Intracellular Ice Growth with Cell-Permeating Small-Molecule Ice Recrystallization Inhibitors. Langmuir, 2019. 35(23): p. 7452–7458.

24. Endo, A., et al., Impact of kestose supplementation on the healthy adult microbiota in in vitro fecal batch cultures. Anaerobe, 2020. 61: p. 102076.

25. Sun, Z., et al., Comprehensive assessment of functional effects of commonly used sweeteners on ex vivo human gut microbiome. bioRxiv, 2022.

