## Supporting information for "An economic and robust TMT labeling approach for high throughput proteomic and metaproteomic analysis"

**Table of Contents:**

Detailed experimental procedures

Figure S1

Figure S2

Figure S3

### Detailed Experimental Procedures

#### Microbiota culturing and sample preparation

The protocol for human stool sample collection (# 20160585-01H) was approved by the Ottawa Health Science Network Research Ethics Board at the Ottawa Hospital, Ottawa, Canada. Stool sample collection, pre-processing and live microbiota bio-banking were performed as described previously [1]. Briefly, fresh stools were collected from healthy adult volunteers, immediately immersed in deoxygenated phosphate-buffered saline (PBS) containing 10% (v/v) glycerol and 0.1% (w/v) L-cysteine hydrochloride, homogenized to make a 20% (w/v) fecal slurry and filtered using sterile gauzes to remove large particles. The processed fecal slurry aliquots were either cultured directly or stored at –80°C prior to further microbiome culturing using the previously established RapidAIM workflow [2].

For the evaluation of TMT labeling efficiency and method development, microbiomes from two individuals (V45 and V46) were harvested, washed using PBS and combined for protein extraction as described previously [3]. Briefly, microbiota cells were re-suspended in lysis buffer containing 4% SDS (w/v), 8 M urea, 50 mM Tris-HCl (pH 8.0), and Roche cOmplete™ mini protease inhibitor. Lysis was carried out with ultrasonication with an amplitude of 50% and a pulse of 10s on/10s off for 10 min using QSonica Q700 sonicator (QSonica LLC., Newtown, USA). After sonication, samples were centrifuged at 16,000 g at 8°C for 10 min to remove any cell debris, and the supernatant was transferred to new tube for precipitation using 5-fold volumes of ice-cold protein precipitation buffer (acetone/ethanol/acetic acid, 50/50/0.1 (v/v/v)) overnight at −20°C. Proteins were then pelleted, washed with acetone, and re-suspended in 6 M urea, 50 mM ammonium bicarbonate buffer for in-solution trypsin digestion. The resulting protein digests were pooled, desalted using C18 column, and eluents were dried and re-suspended in 100 mM TEAB with or without ACN for TMT labeling as described below.

In the ice recrystallization inhibitor (IRI) study, fresh stools were collected from individual V51 and were processed as described above to make 20% (w/v) fecal slurry. The resulting fecal slurry were cultured directly, or were stored at –80°C for two weeks with or without ice recrystallization inhibitors (IRIs). Two different IRIs were tested with five different concentrations each, namely IRIa (β-4-bromophenyl-D-glucose,1.68 mg/ml, 3.35 mg/ml, 5.03 mg/ml, 6.70 mg/ml, 15.08 mg/ml) and IRIb (*N*-2-fluorophenyl-D-gluconamide, 0.29 mg/ml, 0.58 mg/ml, 1.45 mg/ml, 2.89 mg/ml, 5.79 mg/ml). For each condition, the microbiotas were cultured with and without kestose (5 mg/ml) in triplicates for 24 hours, and the cultured microbiotas were harvested and processed as described above for metaproteomic sample preparation and TMT labeling using dry TMT workflow established in this study (Figure 1).

#### Caco-2 cell culture and sample preparation

Human intestinal Caco-2 cells were purchased from the American Type Culture Collection (ATCC) and grown at in Dulbecco's Modified Eagle Medium (DMEM) supplemented with 10% fetal bovine serum at 37°C with 5% CO_2_. Cells used were at passage 15 and grown to 90% confluence. Cells were lifted from plates, washed twice with PBS, and pellets were stored at –80°C until use. Approximately 1.5 × 10^7^ cells were lysed on ice for 5 minutes in 500 μl 1% SDS in 100 mM TEAB buffer containing Thermo Scientific™ Halt™ protease inhibitor cocktail, followed by nuclease treatment using 250 U Pierce™ Universal Nuclease (catalog# 88700) to shear DNA and reduce sample viscosity. The protein content was measured with the BCA assay. Protein disulphides were reduced with 15 mM tris(2-carboxyethyl)phosphine (TCEP) at 55°C for 1 hour then alkylated with 25 mM iodoacetamide for 30 minutes at 21°C protected from light. Protein was precipitated with the addition of 6 volumes of ice-cold acetone, the precipitation proceeded at –20°C overnight. The protein was pelleted by centrifugation at 8000 g for 10 min at 4°C, air-dried briefly, dissolved to 2 μg/μl in 100 mM TEAB followed by digestion with 1: 50 trypsin: protein ratio overnight at 37°C with shaking on thermomixer. After trypsin digestion, the peptide solution was diluted in 100mM TEAB with or without ACN for further TMT labeling as described below.

#### TMT reagents preparation and labeling

TMT11plex label reagents (catalog number A34808) were purchased from ThermoFisher Scientific Inc. For the evaluation of TMT labeling efficiency, equal amount of the 11 channels of TMT reagents were pooled to generate a TMT mix. A portion of the TMT mix was stored in ACN solution at –80°C until use, and the remaining was aliquoted into low-binding tubes with 50 μg or 100 μg per aliquot, completely dried using a CentriVap vacuum concentrator (Labconco Corp.) and stored at –80°C for use. All TMT labelling reactions were performed in triplicates for each group. For the labeling of microbiome-derived samples, 20 μg or 10 μg of desalted peptides were dissolved in 20 μl 100 mM TEAB and mixed with 5 μl ACN containing 100 μg TMT mix for the labeling using the conventional TMT labeling workflow. In the case of dry TMT labeling using 100 μg TMT, 20 μg or 10 μg desalted peptides were dissolved in 25 μl 100 mM TEAB (with 20% ACN) and added directly into the tubes containing 100 μg dry TMT mix for labeling at 25°C for 2 hours. Similarly, in the groups of labeling using 50 μg dry TMT, 12 μg or 6 μg desalted peptides were dissolved in 15 μl 100 mM TEAB (with 20% ACN) were directly added into 50 μg dry TMT mix tube for labeling. For the labeling of Caco-2 cell samples using the conventional workflow, the protein digest solution (20 μg peptides) was directly diluted in 100mM TEAB to 19 μl and mixed with 100 μg TMT mix in 8 μl ACN for labeling at 25°C for 2 hours. For dry TMT labeling, 20 μg or 12 μg peptide digests were diluted into 27 μl or 15 μl peptide solution containing a final concentration of 30% ACN / 100mM TEAB, respectively, and added directly into tubes containing 100 or 50 μg dry TMT for labeling at 25°C for 2 hours. All TMT labeling reactions were quenched using a final concentration of 0.4% hydroxylamine for 15 min following by desalting prior to LC-MSMS measurements.

To prepare dry TMT plates or tube strips, 5 mg TMT11plex label reagents were equilibrated to room temperature, re-suspended in pure ACN to make 1 mg/ml stock solution, and aliquoted into plates or strip tubes with 50 μg reagent in each well/tube. The reagent plates/tube strips were then completely dried using a CentriVap vacuum concentrator (Labconco Corp.) and stored at –80°C prior to use. For convenience, a 96-well plate and a 12-tube strip were used with each row or strip being a full labeling reagent set as indicated in Figure 1 (left to right: 126, 127N, 127C, 128N, 128C, 129N, 129C, 130N, 130C, 131N, 131C, empty). In this study, protein digest of Caco-2 samples were diluted to 0.74 μg/μl in buffer containing 30% ACN / 100mM TEAB. 15 μl of peptide solution (~11 μg) were added directly to each tube of 2 tube-strips for labeling at 25°C for 2 hours, then were quenched by adding 2 μl of 5% hydroxylamine and incubating 15 minutes at room temperature. The individual labels were then combined, lyophilized to dryness and desalted with HyperSep C18 tips (ThermoFisher Scientifc Inc.) according to the manufacturer’s directions. An aliquot of each mixture (100 ug) was further fractionated into 8 fractions with Pierce High pH reversed phase fractionation kit (ThermoFisher Scientifc Inc.) according to the manufacturer’s directions. Desalted and Fractionated samples were lyophilized and dissolved at 500 ng/μl with 0.1% FA for LC-MSMS analysis. For microbiome samples in IRI study, desalted peptides were re-suspended in 100 mM TEAB (with 20% ACN) to make a final concentration of 0.8-1 μg/μl. An aliquot of each sample was combined to generate a pooled sample, which will be used as reference (channel TMT channel 126 in this study for all experiments). Then, 15 μl of the peptide solution or reference sample were added to each well of 96-well plate, mixed well, and incubated at 25°C with agitation for 2 hours. After reaction, 15 μl 0.8% hydroxylamine was added to quench the reaction for 15 min following by acidifying by adding 60 μl of 5% formic acid. Samples from each plate row were then combined to generate a mixture for either desalting prior to LC-MSMS analysis.

#### LC-MSMS analysis

Tryptic peptides derived from microbiome samples were analyzed on an Exploris 480 mass spectrometer (ThermoFisher Scientific Inc.) coupled with an UltiMate 3000 RSLCnano liquid chromatography system. The separation of peptides was performed on an analytical column (75 μm × 50 cm) packed with reverse phase beads (1.9 μm; 120-Å pore size) with 1 or 2-hour gradient from 5 to 35% acetonitrile (v/v) at a flow rate of 300 nl/min. The MS method consisted of a full MS scan from 350 to 1200 m/z at resolution 120,000, followed by data-dependent MS/MS scans with a first mass of 110 m/z, MS2 resolution of 45,000, isolation window of 0.7 m/z, and HCD collision energy of 36%. A dynamic exclusion duration was set to 60 s. All data were recorded by Xcalibur version 4.3 in centroid mode and exported into RAW format for further peptide/protein identification and quantification.

Tryptic peptides derived from Caco-2 cells in this study were analyzed using a Fusion Lumos Tribrid mass spectrometer (ThermoFisher Scientific Inc.) coupled with EASY-nLC™ 1200 System (ThermoFisher Scientific Inc.). For each injection 500 ng peptides were analyzed by loading onto a NanoViper Acclaim pepmap 100 trap column (75 µm × 20 mm with 3 µm beads) and desalting with 0.1% formic acid in water (solvent A) before separating on a EasySpray pepmap C18 reverse-phase analytical column (75 µm × 150 mm with 2 µm beads). Chromatographic separation was achieved at a flow rate of 0.300 µl/min over 100 min in eight linear steps as follows (solvent B was 0.1% formic acid in 80% acetonitrile): initial, 5% B; 80 min, 25% B; 90 min, 40% B; 95 min, 95% B; 100 min, 95% B. The eluting peptides were analyzed in data-dependent mode MSMS. A MS survey scan of 400−1600 m/z was performed in the Orbitrap at a resolution of 120000, auto maximum injection time and an AGC target of 4 x10^5^. The top speed mode was used to select ions for MS2 analysis with dynamic exclusion 20 s with a ± 10 ppm window. During the MS2 analyses precursors were isolated using a width of 0.7 m/z and fragmented by HCD followed by Orbitrap analysis at a resolution of 50000. HCD precursors were fragmented with collision energy of 38% with auto maximum injection time and an AGC target of 1.25 x10^5^.

#### Bioinformatics

The Database searches for the gut microbiome data set were performed using MetaLab (version 2.1) with an integrated human gut microbial gene catalog database (containing 9.9 million microbial protein sequences) and MetaPro-IQ workflow [4-6]. Briefly, a dataset-specific reduced database was generated by matching all MS spectra and the reduced database was used for a second database search using MaxQuant Andromeda algorithm [7]. For TMT labeling efficiency evaluation, TMT (+ 229.163 Da) was specified as variable modifications on lysine and peptide N termini; and for over-labeling evaluation, database search was performed with Reporter ion MS2 (TMT11) mode and, additionally, TMT (+ 229.163 Da) on histidine (H) or serine (S), threonine (T), and tyrosine (Y) were set as variable modifications. For quantitation, the database search was performed with a standard Reporter ion MS2 (TMT11) mode. For all searches, Carbamidomethyl (C) was set as a fixed modification, and Oxidation (M) and Acetyl (Protein N-term) as variable modifications.

Protein and peptide identification of Caco-2 data set was performed using both MaxQuant and Proteome Discoverer 2.4 (ThermoFisher Scientific, Inc.). For both searches, a human SwissProt database (downloaded March 9, 2022, 20307 entries) was used. The same MaxQuant search parameters as described above for the microbiome dataset were used for labeling efficiency check, over labeling check, and quantitation. For the Proteome Discoverer search, the SequestHT algorithm was used with parameters set similar to those used in MaxQuant search. Briefly, all MSMS spectra were searched against human database with 10ppm MS1 and 0.05 Da MS2 tolerances specifying maximum two missed cleavages. Fixed modifications were cysteine carbamidomethyl (+57.021 Da), while variable modifications were peptide N-terminal and lysine TMT (+229.163 Da), methionine oxidation (+15.995 Da), asparagine and glutamine deamidation (+0.984 Da), protein N-terminal acetylation (+42.011 Da), protein N-terminal Met-loss (-131.040 Da), protein N-terminal Met-loss + acetyl (-89.030 Da). The Percolator node was used to improve the rate of peptide identifications using semi-supervised machine learning to discriminate between correct and decoy spectrum identifications.

For the calculation of TMT labeling efficiency, identified PSMs were used and only PSMs with TMT modification on all lysine resides and free peptide N termini were considered as “fully labeled”. PSMs that did not have any TMT modification and has at least one accessible amine were considered as “not labeled”. PSMs that contained at least one TMT modification and were not fully labeled were considered as “partially labeled”. Over-labeling rate was calculated by dividing the number of identified PSMs with TMT modification on either H, S, T or Y by the total number of identified PSMs.

Histograms were generated using *ggplot* in R (version 4.1.2). Box plot of ratio distribution was generated using R package ggpubr and ggboxplot function. Principal component analysis (PCA) and plotting was performed in R as well using R function *autoplot* and *prcomp*.

### Supplementary Figures

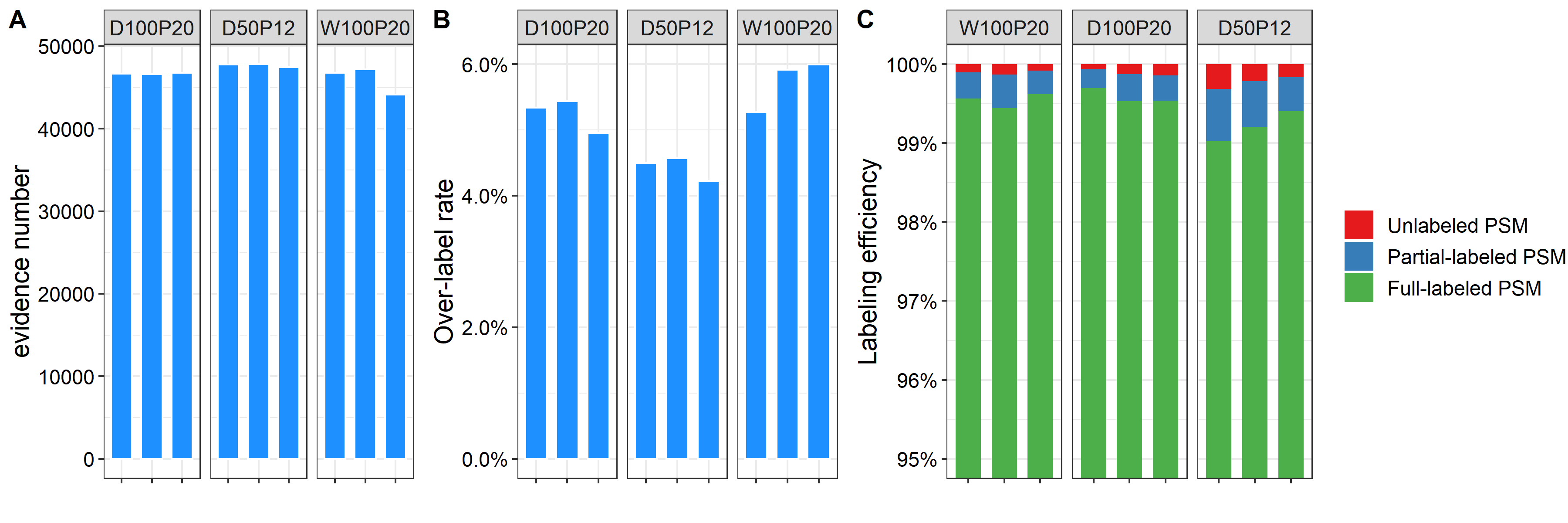

**Figure S1** Comparison of labeling efficiency between convenient dry TMT labeling workflow with standard wet TMT workflow in Caco-2 cell samples. Identified PSM evidence (A), TMT labeling efficiency (B), and over labeling rate (C) were calculated using database search results of Proteome Discoverer.

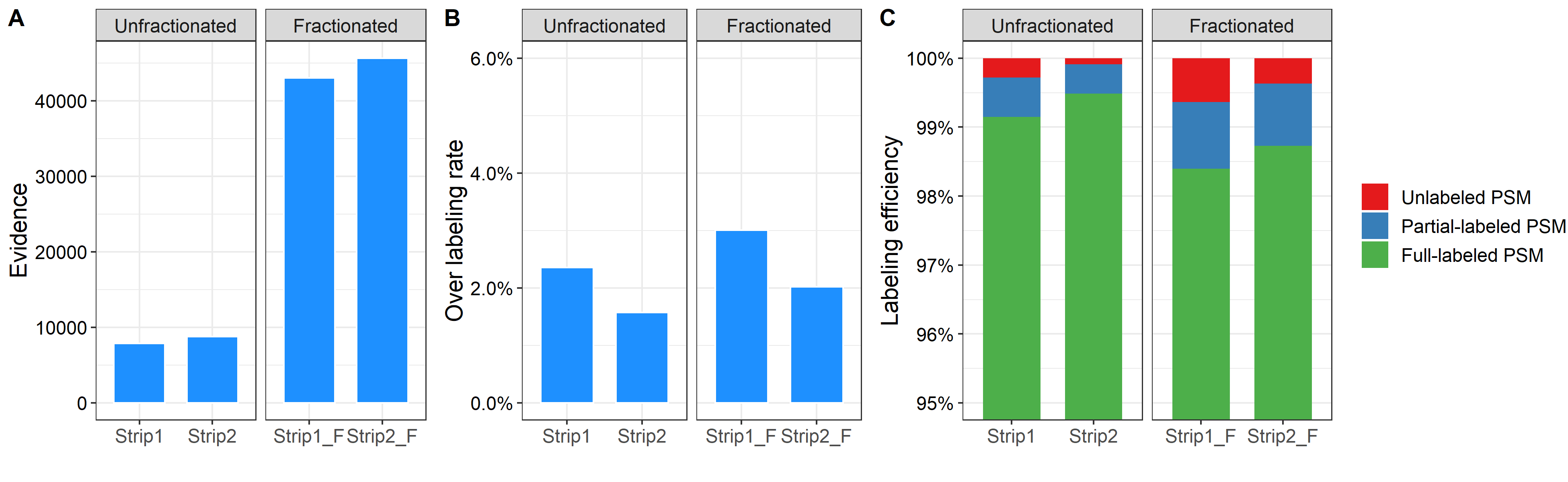

**Figure S2**. Evaluation of labeling efficiency of convenient dry TMT labeling workflow in fractionated and unfractionated Caco-2 samples. (A) Identified PSM evidence; (b) TMT over labeling rate on histidine, serine, threonine, or tyrosine residues; (C) TMT labeling efficiency.

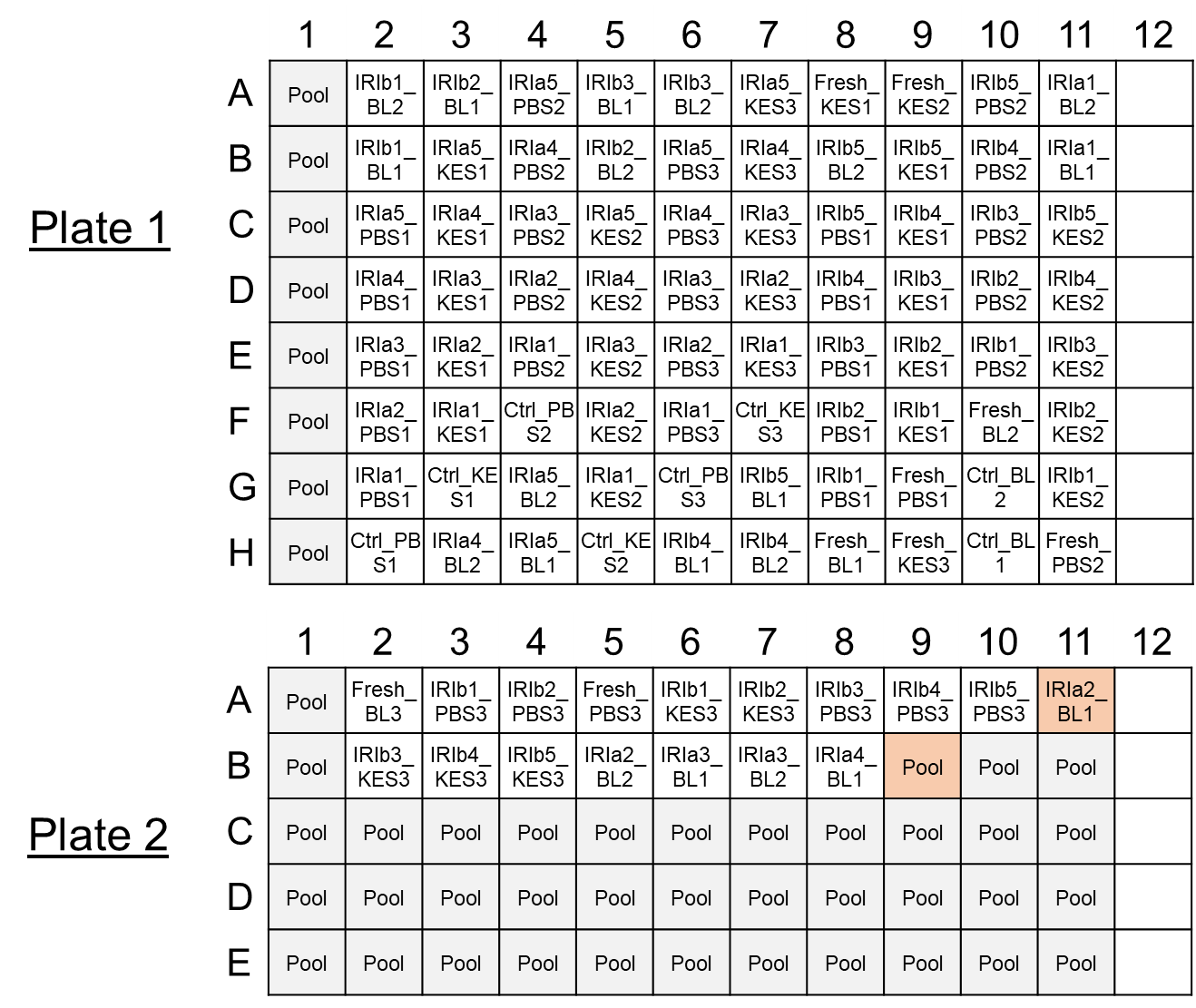

**Figure S3**. Layout of IRI study samples in 96-well TMT plates. The two wells highlighted in pink color in Plate 2 were removed for further analysis due to mislabeling.

**Supplementary References:**

1.	Zhang, X., et al., *Evaluating live microbiota biobanking using an ex vivo microbiome assay and metaproteomics.* Gut Microbes, 2022. **14**(1): p. 2035658.

2.	Li, L., et al., *RapidAIM: a culture- and metaproteomics-based Rapid Assay of Individual Microbiome responses to drugs.* Microbiome, 2020. **8**(1): p. 33.

3.	Zhang, X., et al., *Assessing the impact of protein extraction methods for human gut metaproteomics.* J Proteomics, 2018. **180**: p. 120-127.

4.	Zhang, X., et al., *MetaPro-IQ: a universal metaproteomic approach to studying human and mouse gut microbiota.* Microbiome, 2016. **4**(1): p. 31.

5.	Cheng, K., et al., *MetaLab: an automated pipeline for metaproteomic data analysis.* Microbiome, 2017. **5**(1): p. 157.

6.	Li, L., et al., *iMetaLab Suite: A one‐stop toolset for metaproteomics.* iMeta, 2022: p. e25.

7.	Cox, J., et al., *Andromeda: a peptide search engine integrated into the MaxQuant environment.* J Proteome Res, 2011. **10**(4): p. 1794-805.
